# Multinucleation resets human macrophages for specialized functions at the expense of mononuclear phagocyte identity

**DOI:** 10.1101/2022.08.22.504763

**Authors:** Kourosh Ahmadzadeh, Marie Pereira, Margot Vanoppen, Eline Bernaerts, Jeong-Hun Ko, Tania Mitera, Christy Maksoudian, Bella B Manshian, Stefaan Soenen, Carlos D Rose, Patrick Matthys, Carine Wouters, Jacques Behmoaras

**Affiliations:** Laboratory of Immunobiology, Department Microbiology, Immunology and Transplantation, Rega Institute, KU Leuven – University of Leuven, Leuven, Belgium; Centre for Inflammatory Disease, Department of Immunology and Inflammation, Hammersmith Hospital, Imperial College London, Du Cane Road W12 0NN, London, UK; NanoHealth and Optical Imaging Group, Translational Cell and Tissue Research Unit, Department of Imaging and Pathology, KU Leuven, Herestraat 49, B3000 Leuven, Belgium; Translational Cell and Tissue Research Unit, Department of Imaging and Pathology, KU Leuven, Herestraat 49, B3000 Leuven, Belgium; Division of Pediatric Rheumatology Nemours Children’s Hospital, Thomas Jefferson University, Philadelphia, United States of America; Division Pediatric Rheumatology, UZ Leuven, Leuven, Belgium; European Reference Network for Rare Immunodeficiency, Autoinflammatory and Autoimmune Diseases (RITA) at University Hospital Leuven, Leuven, Belgium; Programme in Cardiovascular and Metabolic Disorders and Centre for Computational Biology, Duke-NUS Medical School Singapore, Singapore

## Abstract

Macrophages undergo plasma membrane fusion and cell multinucleation to form multinucleated giant cells (MGCs) such as osteoclasts in bone, Langhans giant cells (LGCs) as part of granulomas or foreign-body giant cells (FBGCs) in reaction to exogenous material. While osteoclast multinucleation is a prerequisite for vertebrate bone homeostasis, the effector function resulting from LGC and FBGC multinucleation is less well-defined. More generally, how multinucleation *per se* contributes to functional specialization of mature mononuclear macrophages remains poorly understood in humans. Here, we integrated comparative transcriptomics with functional assays in purified mature mononuclear and multinucleated human osteoclasts, LGCs and FBGCs. Strikingly, in all three types of MGCs, multinucleation causes a pronounced down-regulation of mononuclear phagocyte identity. We show enhanced lysosome-mediated intracellular iron homeostasis promoting MGC formation. The transition from mononuclear to multinuclear state is accompanied by cell specialization specific to each polykaryon. Enhanced phagocytic and mitochondrial function associate with FBGCs and osteoclasts, respectively. Moreover, only B7-H3 (CD276)-expressing human LGCs can form granuloma-like clusters *in vitro*, suggesting that LGC multinucleation potentiates T cell activation. These findings demonstrate how cell-cell fusion and multinucleation reset human macrophage identity as part of an advanced maturation step that confers MGC-specific functionality.

## Introduction

Cell-cell fusion in monocyte/macrophage lineage leads to the formation of diverse multinucleated giant cells (MGCs), depending on the tissue microenvironment. MGCs can be classified into osteoclasts, Langhans giant cells (LGCs) and foreign body giant cells (FBGCs), on the basis of their anatomical site, morphology and function during homeostasis or inflammatory disease [1-3]. Osteoclasts are macrophage-derived multinucleated cells specialized in vertebrate bone remodelling and can be considered as homeostatic MGCs that turn-over during adult life [4]. Conversely, LGCs and FBGCs are preferentially found in pathological sites during inflammatory processes. LGCs are the hallmark of infectious (tuberculosis) and non-infectious (sarcoidosis, Blau syndrome) granulomatous diseases [1]. FBGCs can be found at the site of implanted prostheses or medical devices [5] and are specialized in complement-mediated phagocytosis of large particles [6]. To date, the precise role of LGCs remains unclear. Recent studies have provided the immune landscape of infectious granulomas at a single-cell resolution [7, 8] but whether LGC presence is beneficial or detrimental to the disease outcome is currently open to debate.

Despite their divergent cell function, the common feature between osteoclasts, LGCs and FBGCs is their monocyte/macrophage origin and their multinucleated cell appearance following cell-cell fusion. In order to give rise to mature multinucleated cells, macrophages go through two key phases of cell differentiation. The first is governed by tissue-dependent signals that lead to fusion-competency (e.g. RANKL for bone osteoclasts [9]). This step is followed by cell-cell fusion [10] and multinucleation of mature macrophages and formation of MGCs with specialized functions [11].

Among the three types of MGCs, studies on osteoclasts improved considerably our understanding of how lineage-specific signals followed by cell-cell fusion shape cell activity during homeostasis and disease. Indeed, functional osteoclasts capable of resorbing bone are multinucleated macrophages that differentiate through the concerted action of macrophage-colony-stimulating factor (M-CSF) and RANKL [9, 12-14]. Osteoclasts originate from embryonic erythro-myeloid progenitors and their postnatal maintenance is mediated by acquisition of new nuclei from circulating blood cells [15]. Once mature, multinucleated osteoclasts can undergo fission and form transcriptionally distinct cells called osteomorphs [16]. Mature osteoclasts contain up to approximately 20 nuclei in normal human bones [17, 18] and during pathological conditions, active osteoclasts may contain over 50 nuclei [19], indicative of an association between the number of nuclei and cell activity. From a genetic point of view, conserved transcriptional gene networks in multinucleating mature osteoclasts control their resorption activity and the resulting bone mass [3, 20, 21]. In summary, fusion and multinucleation of osteoclasts give rise to a separate, specialized cell stage, which is absent in mononuclear cells. Paradoxically, even though extensive efforts went into understanding the biology of osteoclast differentiation (i.e. osteoclastogenesis), relatively less is known about the transition from a RANKL-induced mononuclear into a fused multinuclear osteoclast state and the resulting post-fusion transcriptional reprogramming in these cells.

The molecular consequences of cell-cell fusion and multinucleation remain poorly defined in mature human osteoclasts, LGCs and FBGCs. One obstacle for side-by-side comparison of fusing MGCs has been the identification of lineage determining factors and their respective efficacy to confer fusion-competency. RANKL is a well-established *in vitro* differentiation signal for osteoclasts but the soluble factors responsible for triggering cell fusion and multinucleation in LGCs and FBGCs are relatively less studied. Although previous evidence points towards IFN-γ and IL-4 in LGCs and FBGCs respectively [22, 23], it is not understood to what extend these two stimuli recap the functional characteristics of LGCs and FBGCs in humans. Here we investigated the mechanisms governing macrophage fusion and multinucleation in mature LGCs, FBGCs and osteoclasts differentiated from healthy donors. We first showed that the usage of a single stimulus (RANKL for osteoclasts, IFN-γ for LGCs and IL-4 for FBGCs) in presence of M-CSF, can recap the typical morphological appearance of these primary human cells. The isolation of purified mononuclear and multinucleated LGCs, FBGCs and osteoclasts allowed us to perform comparative transcriptomics and focus on shared and cell-type specific observations during macrophage multinucleation. As part of shared pathways, we show that multinucleation causes a drastic down-regulation of macrophage identity meanwhile lysosome-dependent iron homeostasis is enhanced. Cell type-specific features include FBGCs showing improved ability of phagocytosis and osteoclasts maximizing their mitochondrial activity following multinucleation. We demonstrate that B7-H3-expressing human LGCs can form granuloma-like clusters *in vitro*. Our results show that macrophage fusion and multinucleation reprograms the cell for specialized functions at the expense of a loss of core mononuclear phagocyte signature.

## RESULTS

### Fusion and multinucleation reshapes the myeloid transcriptome in differentiated human MGCs

We generated three types of multinucleated giant cells (MGCs) using macrophages isolated from human donor-derived peripheral blood mononuclear cells (PBMCs) (**Figure 1A**). In addition to RANKL used for osteoclasts, IFN-γ and IL-4 were added to generate mature LGCs and FBGCs, respectively (**Figure 1A**). As expected, IFN-γ induced LGCs were characterized by a ring of nuclei along the cell border, while IL-4-induced FBGCs showed scattered nuclei throughout the cytoplasm (**Figure 1A** and **B**) [22, 24, 25]. The distinct morphological appearance of these three types of MGCs (**Figure 1B**) suggested cell type-specific functional properties and shared mechanisms underlying macrophage multinucleation. Macrophage fusion and multinucleation include a differentiation step from progenitor cells that is achieved with lineage-specific signals (e.g. RANKL, IFN-γ, IL-4). Following the differentiation step, cell fusion and multinucleation lead to mature LGCs, FBGCs and osteoclasts [1, 3]. Here we aimed to investigate the functional consequences of human macrophage fusion/multinucleation post-cell differentiation. We first confirmed that MGCs are generated through cell membrane fusion in all 3 cell types (**Supplementary Figure 1A**). Next, we set-up a cell sorting strategy (**Supplementary Figure 1B, C and methods**), resulting in >85% purity in mononuclear and multinucleated LGCs, FBGCs and osteoclasts after their respective differentiation with lineage-specific soluble factors (**Figure 1B**). RNA-seq analysis between mononuclear and multinucleated cells allowed us to minimize cofounding lineage signal effects (RANKL, IFN-γ, IL-4) and focus on transcriptional pathways arising from multinucleation *per se*. The top differentially expressed genes between mononuclear and multinucleated cells included markers of tissue-resident macrophages (*LYZ* [26], *MSA4A7* [27]) and of pattern recognition receptors (*TLR2, CLEC4E, PTAFR, FCN1*) (**Figure 1C**), suggesting a transcriptional reprogramming of myeloid identity, which we set out to investigate in more detail.

**Figure 1.**
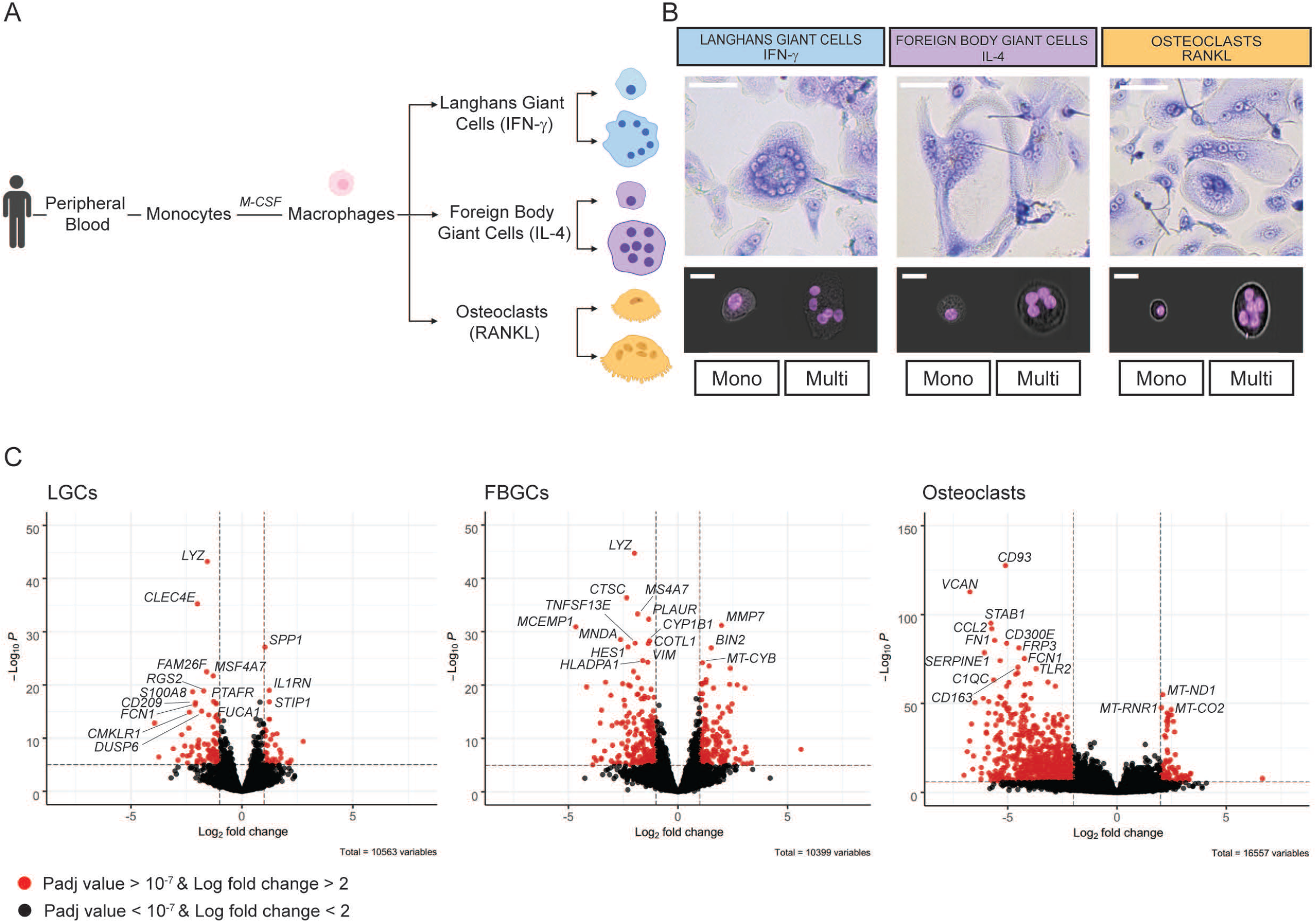
Human macrophage fusion and multinucleation reprograms the transcriptome in differentiated Langhans giant cells (LGCs), foreign body giant cells (FBGCs) and osteoclasts. **(A)** Schematic overview of the PBMC-derived generation of differentiated mononuclear and multinucleated LGCs, FBGCs and osteoclasts using IFN-γ, IL-4 and RANKL, respectively. **(B)** Light microscopy images of LGCs, FBGCs and osteoclasts stained with Giemsa staining (upper panel). Representative Hoescht dye images acquired by ImageStream (lower panel) of sorted cells showing mononuclear (mono) and multinucleated (multi) populations for each cell type. **(C)** RNA-seq volcano plots highlighting the top 15 differentially expressed genes between mononuclear and multinucleated populations for each cell type. scale bar,100 µm; ImageStream shown at 20X magnification to visualise the sorting purity.

### Multinucleation suppresses a shared mononuclear phagocyte gene signature in humans

We first analysed the commonly down-regulated genes as a result of multinucleation in LGCs, FBGCs and osteoclasts (**Figure 2A**). These 191 transcripts (**Supplementary Table 1**) belong to pathways that include tuberculosis, phagosome, osteoclasts and more generally the immune system (**Figure 2B**). Remarkably, these commonly down-regulated genes contain a set of transcripts that defines mononuclear phagocytes and we confirmed their multinucleation-induced suppression by qRT-PCR (**Supplementary Figure 1D**). Among these 191 transcripts, phagocytosis/apoptotic cell clearance (*FCGR3A, FCGR2A, HCK, LYN, C1QA, C1QB C1QC*), antigen presentation (*HLA*-*DPA1, HLA*-*DRA, HLA*-*DRB1, HLA*-*DPB1, CD74*), tissue resident macrophage (*CSF1R, MRC1, CD163, MAFB*) transcripts form a protein-protein interaction network (PPI, **Supplementary Figure 2A**). Osteoclast lineage determinant transcription factor *FOS* [28], macrophage-specific transcription factor *MAFB* [29, 30] and LGC multinucleation inducer *TLR2* [31] are among the transcripts forming the core mononuclear phagocyte signature that is suppressed with multinucleation in differentiated LGCs, FBGCs and osteoclasts.

**Figure 2.**
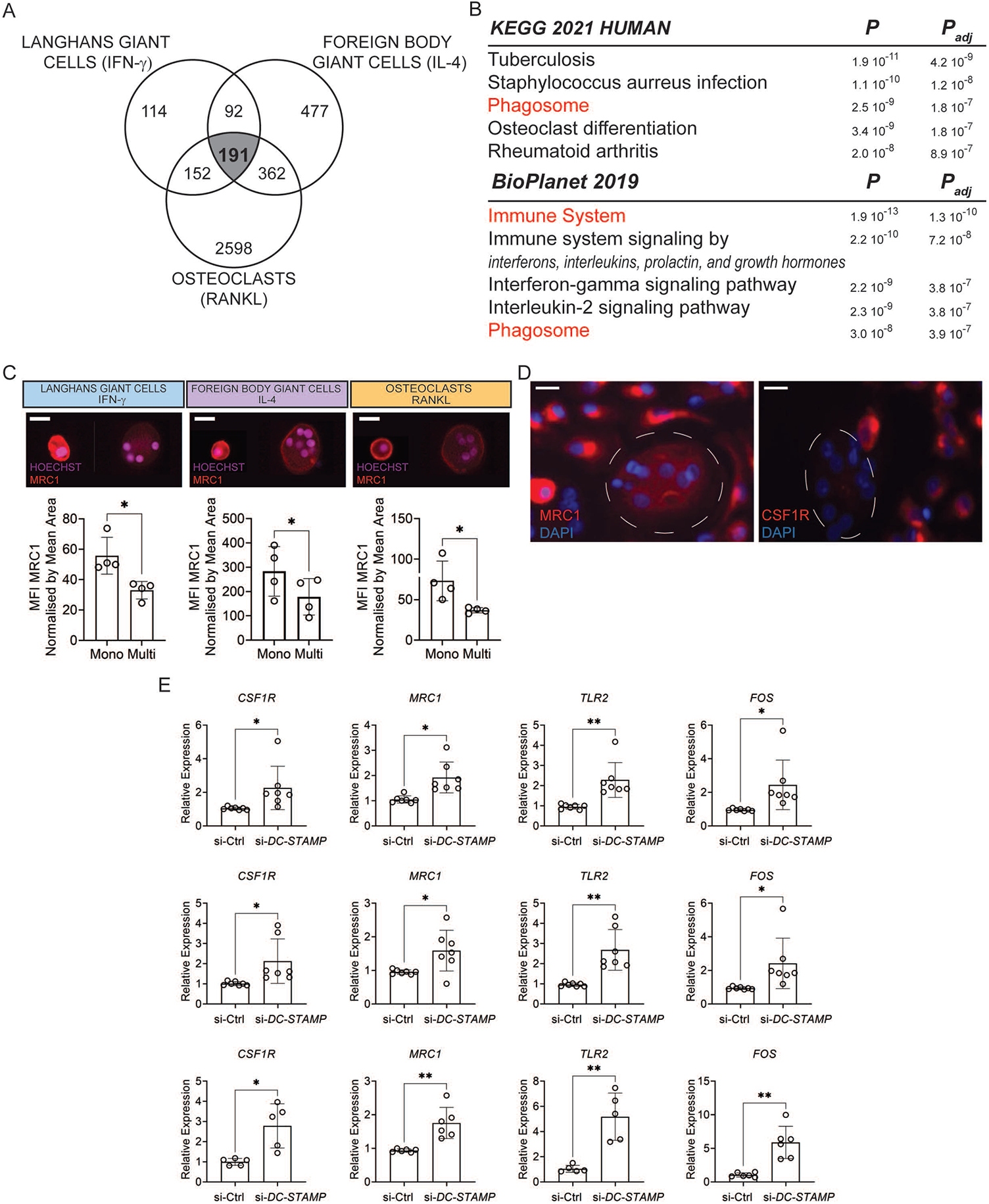
Fusion and multinucleation causes the down-regulation of a shared mononuclear phagocyte gene signature between the LGCs, FBGCs and osteoclasts. **(A)** Venn diagram showing the commonly downregulated transcripts (n=191) as a result of multinucleation in LGCs, FBGCs and osteoclasts. **(B)** KEGG and BioPlanet 2019 pathway analyses on the 191 commonly downregulated genes. Pathways in red are shared between KEGG and BioPlanet. **(C)** MRC1 surface marker expression (red) acquired by ImageStream in mononuclear (mono) and multinucleated (multi) LGCs, FBGCs and osteoclasts stained for Hoechst. Bar graphs (lower panel) represent normalized MRC1 mean fluorescence intensity (MFI), measured by ImageStream; n=4 donors. **(D)** Representative MRC1 (left panel) and CSF1R (right panel) immunofluorescence in mononuclear and multinucleated osteoclasts stained for DAPI (blue). Note the dim MRC1 and CSF1R in multinucleated osteoclasts (dashed lines) compared to surrounding mononucleated ones. **(E)** *CSF1R, MRC1, TLR2* and *FOS* relative expression measured by qRT-PCR for LGCs (upper), FBGCs (middle) and osteoclasts (lower), following the cell membrane fusion regulator DC-STAMP knockdown. si-Ctrl, scrambled siRNA; si-DC-STAMP, DC-STAMP siRNA; n=7 donors. Error bars are mean ± SD; significance tested by paired t-test; *, P < 0.05; **, P < 0.01. Scale bars, 20 µm **(C)** and 100 µm **(D)**.

To validate this fusion-induced suppression of mononuclear phagocyte transcriptome at the protein level, we next focused on two major markers of tissue macrophages: CSF1R [32, 33] and MRC1 (CD206) [34]. Surface expression of MRC1 and CSF1R was measured by ImageStream and immunofluorescence (**Figure 2C** and **D**). To assess cell surface MRC1 levels, an ImageStream gating strategy was applied to mononuclear and multinucleated cells (**Supplementary Figure 2B** and **C**). In accordance with the transcriptomic data, multinucleation significantly suppressed the protein levels of both tissue macrophage markers (**Figure 2C, D** and **Supplementary Figure 2D**).

Multinucleation-induced suppression of a core mononuclear phagocyte signature suggests that cell-cell fusion but not lineage-determinants factors are responsible for the loss of macrophage identity. To strengthen this observation, we tested whether direct disruption of cell fusion can recapitulate the expression changes in *MRC1, CSF1R, TLR2* and *FOS* in multinucleating LGCs, FBGCs and osteoclasts. We thus performed transient RNAi for *DC*-*STAMP*, a plasma membrane regulator of cell fusion in MGCs [21, 35, 36] and obtained >90% knockdown throughout the three primary cell-types (**Supplementary Figure 2E**). Strikingly, *DC*-*STAMP* knockdown reversed the expression of *MRC1, CSF1R, TLR2* and *FOS* in LGCs, FBGCs and osteoclasts (**Figure 2E**), confirming that cell fusion and multinucleation trigger the extinction of mononuclear phagocyte gene signature.

### Lysosome-dependent iron homeostasis drives human macrophage multinucleation

We next investigated the commonly upregulated genes as a result of multinucleation in LGCs, FBGCs and osteoclasts (**Figure 3A**). These 66 gene transcripts (**Supplementary Table 2**) belong to pathways that include lysosome, iron uptake and transport (**Figure 3B**). Optimal lysosomal function is a prerequisite for osteoclast function [37] and iron regulates a broad range of macrophage effector functions, including mitochondrial activity and ATP generation [38]. Specifically, the up-regulated genes in the three cell types include *ATP6V1H, ATP6V1D, TFRC* and *SLC11A2*. These genes encode for the two subunits of the V1 domain of v-ATPase responsible for ATP-dependent lysosomal acidification (*ATP6V1H* and *ATP6V1D*), transferrin receptor (*TFRC*) and endosomal iron transporter (SLC11A2, also known as NRAM2). Recent evidence suggests that when vacuolar or lysosomal acidification is perturbed, this creates a cytoplasmic iron deficiency and defects in mitochondrial metabolism [39-41]. We thus hypothesized that macrophage multinucleation requires an enhanced lysosomal biogenesis that maintains cytoplasmic iron pools needed to sustain the metabolically demanding process of plasma membrane fusion (**Figure 3C** and **D**). To test this hypothesis, we induced lysosomal dysfunction through 4 different pharmacological means. Ammonium Chloride (NH_4_Cl) and hydroxychloroquine Sulfate (Hydroxy S) are pH-disrupting lysosomotropic agents while Bafilomycin A1 (Baf A1) and Concanamycin A (Con A) are vATPase inhibitors [42, 43]. The use of all four inhibitors of lysosomal function caused a decrease in macrophage fusion in LGCs, FBGCs and osteoclasts (**Figure 3E**). Importantly, iron replenishment through plasma membrane but not lysosomal compartment (FeCl_3_ addition) partially rescued the defect in macrophage fusion resulting from lysosomal dysfunction (**Figure 3E**).

**Figure 3.**
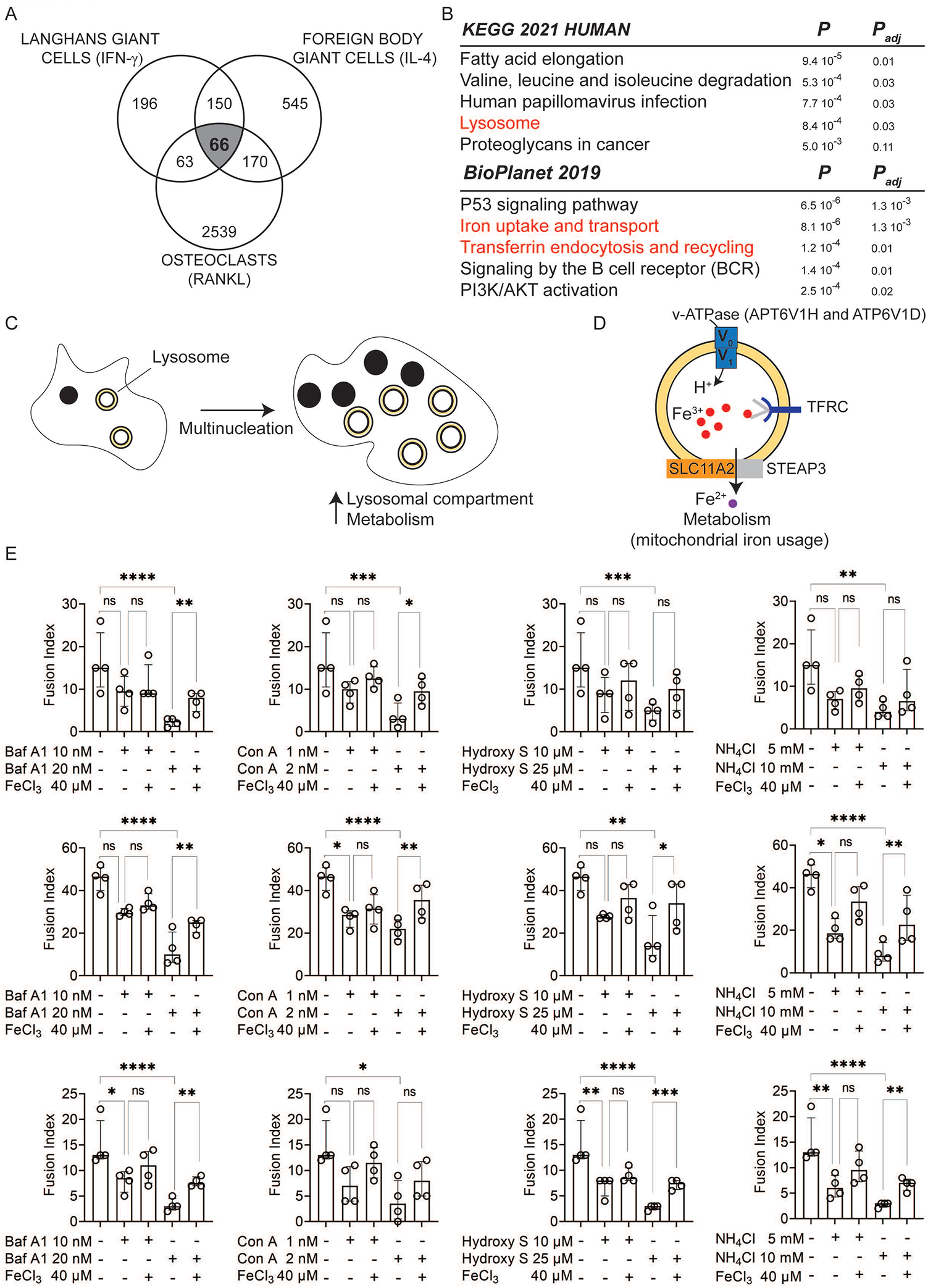
Acidic lysosomal pH upstream iron homeostasis is necessary for macrophage fusion **(A)** Venn diagram showing the commonly upregulated transcripts (n=66) in LGCs, FBGCs and osteoclasts. **(B)** KEGG and BioPlanet 2019 pathway analyses on the 66 commonly upregulated genes. Pathways in red are connected. **(C)** Macrophage fusion and multinucleation causes an increased lysosomal compartment and cell metabolism. **(D)** The commonly up-regulated genes in LGCs, FBGCs and osteoclasts (*ATP6V1H, ATP6V1D, TFRC* and *SLC11A2* and their schematic role in the lysosome-iron pathway. **(E)** The effect of lysosomotropic agents (hydroxychloroquine, hydroxy; ammonium chloride, NH_4_Cl) and v-ATPase inhibitors (Bafilomycin A1, Baf A1; Concanamycin A, Con A) on fusion and multinucleation in LGCs (upper panel), FBGCs (middle panel) and osteoclasts (lower panel). To test the effect of lysosome dysfunction on cellular iron, FeCl_3_ was supplemented. Error bars are median with interquartile range; significance tested by one-way Anova followed by Sídák’s multiple comparisons on log transformed data; n = 4 donors; *, P < 0.05; **, P < 0.01; *** P < 0.001; ****, P < 0.0001.

### B7-H3-expressing human LGCs form granuloma-like clusters

The results above show that multinucleation causes a suppressed mononuclear phagocyte gene signature while lysosomal function upstream iron homeostasis is induced as a result of cell-cell fusion in LGCs, FBGCs and osteoclasts. Having established these shared mechanisms in cell fusion between the three types of multinucleated macrophages, we next investigated cell type-specific pathways. The uniquely differentially expressed transcripts in LGCs (**Figure 4A, Supplementary Table 3**) were enriched for antigen presentation, and adaptive immune system pathways (**Figure 4B**) which are prominent features of infectious granulomas [25]. We found that genes belonging to lipid and atherosclerosis pathway also characterize LGCs (**Figure 4B**) and these were recently linked to LGC formation [44]. Given the multicellular environment that characterizes infectious granulomas and the resulting adaptive immune responses, we reasoned that LGCs, but not FBGCs and osteoclasts, can be organized in granuloma-like clusters *in vitro*. In order to test this hypothesis, donor-derived total PBMCs instead of monocytes were stimulated with either IFN-γ or IL-4 or RANKL. At day 7 of differentiation of monocyte-lymphocyte co-cultures, MGCs were stained for immunofluorescence analysis. Importantly, granuloma-like clusters were formed in LGC culture conditions, and were absent in FBGCs and osteoclasts cultures (**Figure 4C**). These LGC-specific granuloma like clusters contained CD3+ T cells, which were absent in FBGCs and osteoclasts (**Figure 4D** and **Supplementary Figure 3A**), suggesting that these were only retained in IFN-γ containing LGC culture conditions. To further strengthen LGC-T cell link, we focused on the LGC-expressed transcript encoding B7-H3 (also known as CD276), a member of the human B7 family and type I transmembrane protein necessary for T cell activation and IFN-γ production [45]. Although *CD276* transcript levels are upregulated in fusing LGCs and osteoclasts (**Supplementary Table 4**), we found that multinucleation of LGCs but not FBGCs and osteoclasts increased significantly the surface expression of B7-H3 (**Figure 4E** and **Supplementary Figure 3**). In summary, IFN-γ-induced macrophage multinucleation leads to B7-H3 expressing LGCs which are associated with CD3+ T cells and lead to the formation of granuloma-like cell clusters.

**Figure 4.**
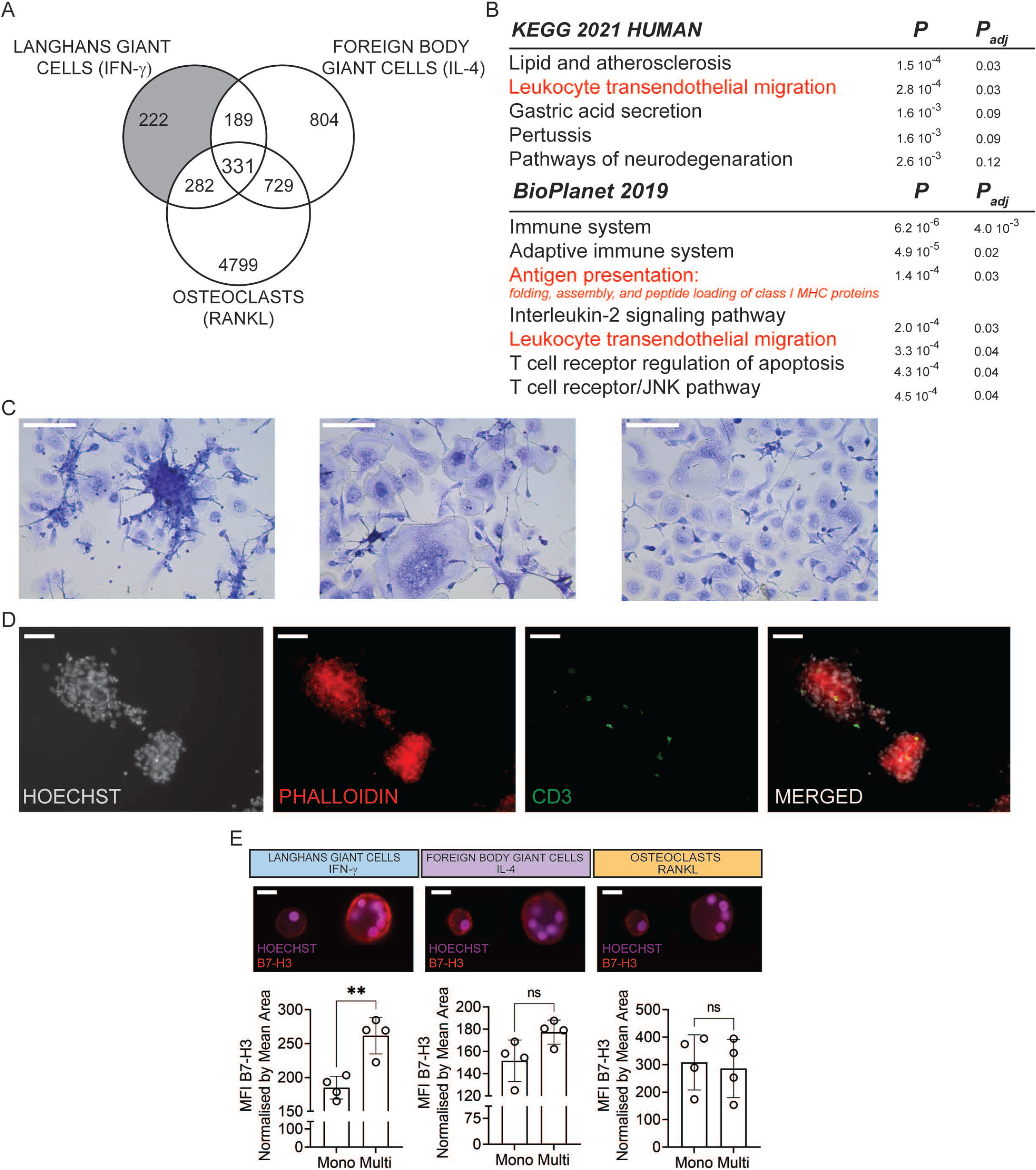
LGCs form in granuloma-like clusters and show increased membrane expression of B7-H3. **(A)** Venn diagram showing the uniquely differentially expressed transcripts in LGCs (grey box). **(B)** KEGG and BioPlanet 2019 pathway analyses on the differentially expressed transcripts in LGCs only. Pathways in red are in common in KEGG and BioPlanet or relevant for LGC function. **(C)** Giemsa stained LGCs (left), FBGCs (middle) and osteoclasts (right) differentiated from PBMCs. Note the cell cluster formation in LGCs only. **(D)** CD3 immunofluorescence (green) in PBMC-derived LGCs. Hoechst (grey) and phalloidin (red) staining show the nuclei and cytoskeleton, respectively. Merged (CD3, Hoechst, phalloidin) image shows the existence of CD3+ T cells in the granuloma-like clusters. **(E)** B7-H3 surface marker expression (red) acquired by ImageStream in mononuclear and multinucleated LGCs, FBGCs and osteoclasts stained for Hoechst. Bar graphs (lower panel) show normalized B7-H3 mean fluorescence intensity (MFI), measured by ImageStream; n=4 donors. Error bars are mean ± SD; significance tested by paired t-test; **, P < 0.01; ns, non-significant. Scale bars, 100 µm **(C and D)** and 20 µm **(E)**.

### FBGCs show improved ability of phagocytosis while osteoclasts maximize their mitochondrial activity following multinucleation

We next focused on FBGC-specific transcripts (**Figure 5A** and **Supplementary Table 5**) and interrogated whether they are indicative of a distinct functional adaptation in these multinucleated cells. Unsurprisingly, FBGCs were enriched for IL-4 signalling pathway together with VEGF-associated genes [46] (**Figure 5B**). Furthermore, pathways like shigellosis which overlap with the regulation of actin cytoskeleton were suggestive of specialized phagocytic activity of FBGCs [6]. We thus quantified phagocytosis of *S. Aureus* -coated beads in mononuclear and multinucleated human LGCs, FBGCs and osteoclasts and showed that multinucleation significantly induced phagocytosis when normalized to cell size in FBGCs but not in the other MGCs (**Figure 5C** and **Supplementary Figure 4**). Eventually, osteoclasts showed a transcriptomic signature robustly associated with mitochondrial activity (i.e. oxidative phosphorylation, TCA cycle and respiratory electron transport) (**Figure 6A** and **B, Supplementary Table 6**). These ATP-generating pathways are indispensable for osteoclasts’ unique and energy-consuming bone resorptive function, which we confirmed in TRAP+ cells (**Figure 6C**). Notably, among the TRAP+ LGCs and FBGCs, only osteoclasts showed hydroxyapatite resorption (**Figure 6C**). To further confirm that multinucleation boosts oxidative phosphorylation in osteoclasts, we quantified oxygen consumption rate (OCAR) and extracellular acidification rate (ECAR) in mononuclear and multinucleated osteoclasts and observed that the spare respiratory capacity and ATP production but not glycolysis measurements are increased significantly with multinucleation (**Figure 6D and E**). Furthermore, osteoclast mitochondria-encoded transcripts were among the genes that showed the largest fold-change upon multinucleation (**Figure 1C**). The multinucleation-induced increase in *MT*-*ND1, MT*-*CYTB, MT*-*CO2, MT*-*CO3* expression was not due to increased mitochondrial copy number as normalized expression levels show significant upregulation in multinucleated vs. mononuclear osteoclasts (**Figure 6F** and **G**). Thus, multinucleation induces maximal respiration in osteoclasts, confirming the unique transcriptional signature in mitochondria-related pathways such as oxidative phosphorylation/TCA cycle and respiratory electron transport chain. In summary, macrophage multinucleation causes a shared down-regulation of mononuclear phagocyte identity while the lysosome-iron pathway is induced (**Figure 6H**). Besides these shared pathways, each polykaryon acquires specialized functions following multinucleation (**Figure 6H**).

**Figure 5.**
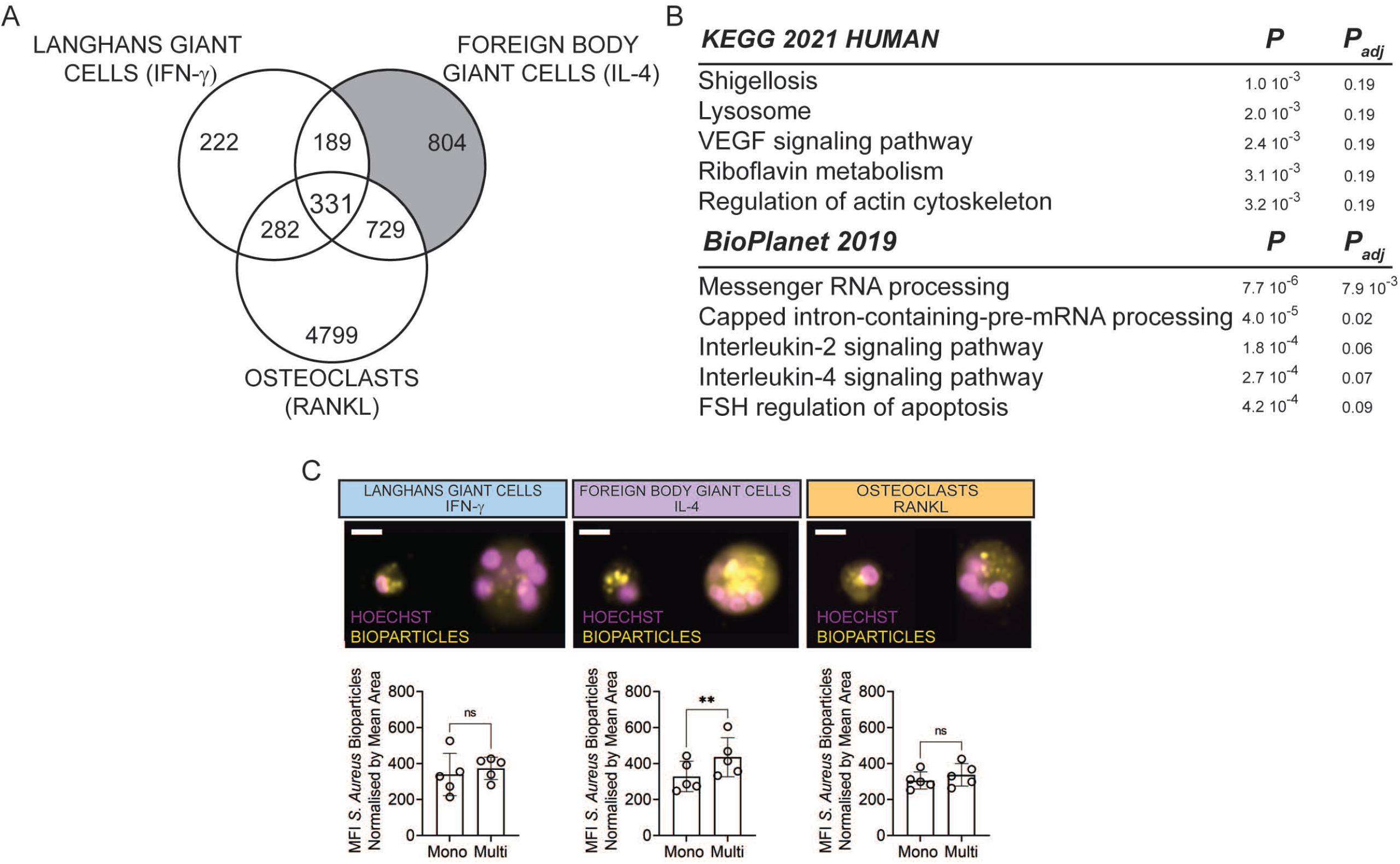
FBGCs show a distinctively enhanced phagocytic capacity. **(A)** Venn diagram showing uniquely differentially expressed transcripts in FBGCs (grey box) **(B)** KEGG and BioPlanet 2019 pathway analyses on differentially expressed transcripts in FBGCs only. **(C)** Phagocytosis quantified by ImageStream analysis in LGCs, FBGCs and osteoclasts. Representative images showing the uptake of *S. Aureus* bioparticles (yellow) in mononuclear (mono) and multinucleated (multi) cells (20X magnification, scale bar = 20 µm). Bar graphs (lower panel) show normalized *S. Aureus* bioparticles mean fluorescence intensity (MFI) measured by ImageStream; n= 5 donors. Error bars are mean ± SD; significance tested by paired t-test; **, P < 0.01; ns, non-significant. Scale bars, 20 µm **(C)**.

**Figure 6.**
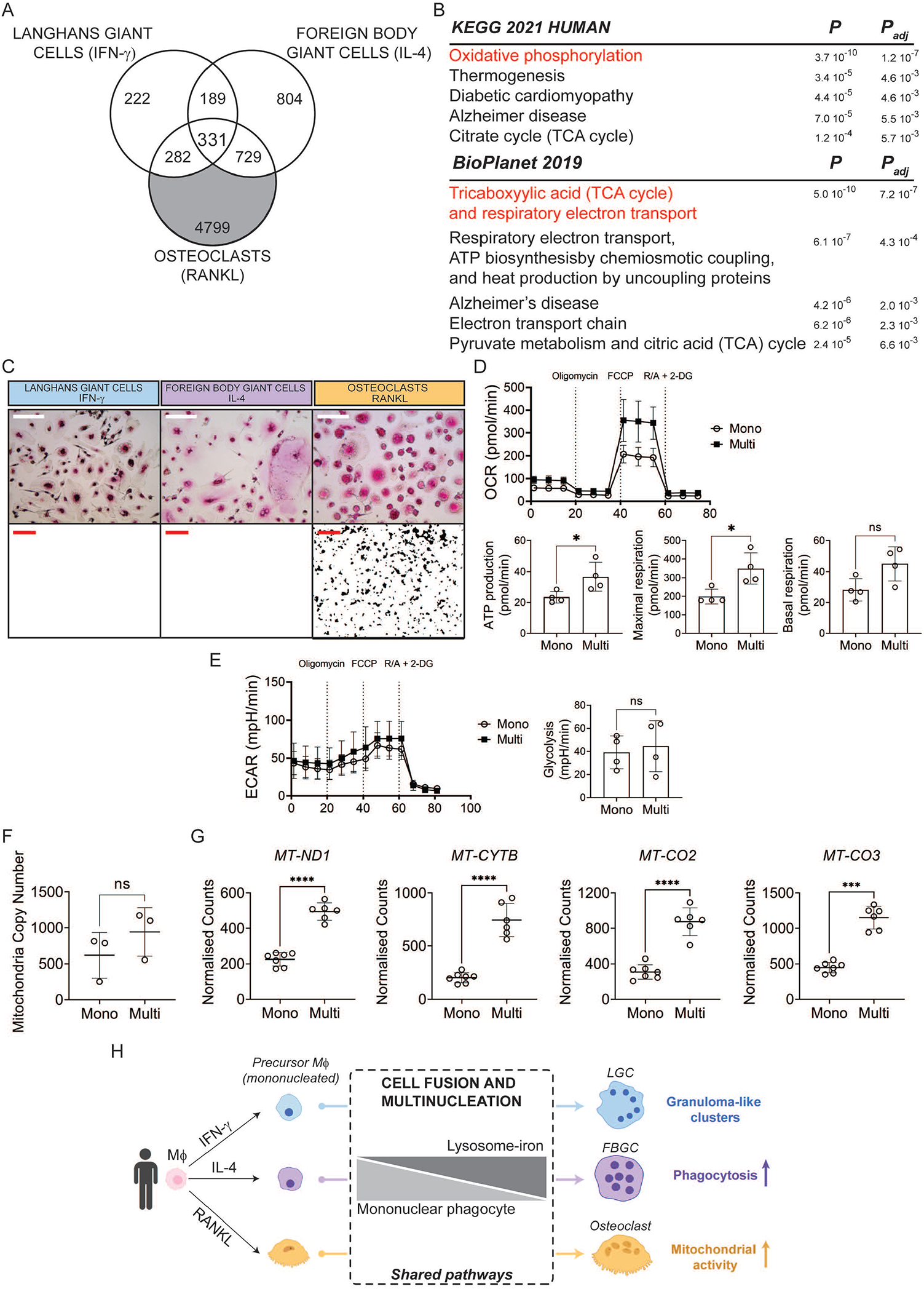
Osteoclasts show a distinctively increased mitochondrial activity. **(A)** Venn diagram showing uniquely differentially expressed transcripts in osteoclasts (grey box). **(B)** KEGG and BioPlanet 2019 pathway analyses on differentially expressed transcripts in osteoclasts only. Pathways in red overlap between KEGG and BioPlanet. **(C)** TRAP immunohistochemistry (upper panel) and hydroxyapatite resorption (lower panel) in LGCs, FBGCs and osteoclasts. Note the presence of TRAP+ MGCs in all cell types and hydroxyapatite resorption only in osteoclasts. **(D)** Oxygen consumption rate (OCR, upper panel), its related readout (middle panel) and extracellular acidification rate (ECAR, lower panel) measured by extracellular flux analysis in sorted mononuclear and multinucleated osteoclasts; n = 4 donors. **(E)** Extracellular acidification rate (ECAR, left) and glycolysis (right) in mononuclear and multinucleated osteoclasts; n = 4. **(F)** Mitochondria copy number measured by qPCR in mononuclear and multinucleated osteoclasts; n=3 donors. **(G)** Relative expression of mitochondrial transcripts in sorted mononuclear and multinucleated osteoclasts after normalisation by mitochondrial copy number. **(H)** Cartoon illustrating that macrophage multinucleation causes a shared down-regulation of mononuclear phagocyte identity while the lysosome-iron pathway is induced. At least n=6 donors; error bars are mean ± SD; significance tested by paired t-test; ***, P<0.001; ****, P < 0.0001; ns, non-significant. Scale bars, 40µm **(C**, upper panel) and 1mm **(C**, lower panel).

## Discussion

Understanding the consequences of cell-cell fusion in terms of macrophage effector functions have been challenging. A multinucleation-induced ‘gain of function’ is a theory put forward to explain why osteoclasts, FBGCs and most likely MGCs of adipose tissue are multinucleated [47]. This hypothesis specifies that multinucleated cell state reprograms the cell for a specific effector function that is not present or is less effective in the mononuclear cell state. Supporting evidence for this concept is the increased lysosome numbers and plasma membrane surface in fused cells, allowing proficient degradative and phagocytic activities, respectively. Indeed, osteoclast multinucleation correlates with the resorptive activity of the cell and regulates bone mass [21]. Similarly, MGCs of the adipose tissue (or adipoclasts) can phagocytose lipid remnants more efficiently when they are multinucleated [48], a feature that is shared with FBGCs which are also specialised in the uptake of relatively large particles [6]. However, these findings do not explain why LGCs are multinucleated in granulomas as these MGCs have been described to allow persistence to *mycobacterium* rather than exerting bactericidal activities [49]. Regarding osteoclasts, although fusion confers resorptive function, recent evidence points toward a previously unappreciated plasticity of mature osteoclasts during inflammation [50, 51]. Hence the functional advantage of macrophage fusion and the underlying pathways is complex in osteoclasts and remain incompletely understood in LGCs and FBGCs.

One reason for this gap in knowledge is the lack of consensus for determining culturing conditions involving LGCs and FBGCs *in vitro*. Furthermore, the reliable isolation of pure populations of mononuclear and multinucleated mature macrophages has hindered the side-by-side comparison of MGC post-fusion events in osteoclasts, LGCs and FBGCs in humans. Here, we show that macrophage cell-cell fusion and multinucleation triggers suppression of a shared mononuclear phagocyte transcriptional signature alongside activation of MGC-specific pathways in human osteoclasts, LGCs and FBGCs. These results were achieved by (i) selecting the lineage-determinant factor representative of *in vivo* morphology of each polykaryon and a (ii) a side-by side comparison of human mononuclear and multinuclear macrophages in terms of their transcriptional and functional reprogramming.

TNF and TNF receptor super-families include RANKL, RANK and osteoprotegerin, and are experimentally proven regulators of osteoclast formation *in vitro* and *in vivo* [9, 52]. On the other hand, IFN-γ is a T-helper cell type 1 (Th1) cytokine, previously demonstrated to be an efficient inducer of LGCs from human monocytes [23, 53]. *In vivo*, IFN-γ is found in human tuberculosis and leprosy granulomas [7, 54] and in non-infectious granulomatous disease such as sarcoidosis [55, 56] and Blau syndrome [57]. All these conditions are associated with the presence of LGCs and T cells (either Th1 or Th17) within the granulomas. Human subjects with defects in the IFN-γ signalling pathway develop mycobacterial infections without formation of granulomas, demonstrating the need of IFN-γ signalling for LGC-containing granuloma formation *in vivo* [58]. Similarly, IL-4 has been shown to induce FBGCs from human monocytes *in vitro* and *in vivo* [22, 59, 60] even though single cell transcriptomics analyses of foreign body reactions have shown that the immune cell landscape may depend on whether the composition of the biomaterial is based on biological matrix or is synthetic [61], the latter being associated with a prominent role of IL-17 [62]. Thus, both IFN-γ and IL-4 have been shown to promote MGC formation *in vitro* and the two cytokines show niche-specific presence in human tissues during inflammatory disease involving MGCs. While macrophage polarization as a result of IFN-γ or IL-4 stimulation is well-documented [63, 64], our results show a robust additional cell reprogramming triggered by multinucleation, suggesting that this advanced maturation step is essential for the cell-specific activity of LGCs and FBGCs *in vivo*.

By choosing a relatively simple cell culture setup involving M-CSF and either IFN-γ, IL-4 or RANKL, we obtained MGCs that served us as a proxy for studying and comparing side-by-side multinucleation of LGCs, FBGCs and osteoclasts. Importantly, both the transcriptomics and functional assays between mononuclear and multinucleated macrophages confirm the key characteristics of each cell type: granuloma-like cluster formation (LGCs), enhanced phagocytosis (FBGCs), heightened mitochondrial activity that fuels bone resorption (osteoclasts). These MGC-specific cell activities together with cell-specific morphological features (e.g. ring-shaped nuclei in LGCs) validate the usage of IFN-γ and IL-4 for the *in vitro* generation of human LGCs and FBGCs, respectively. Our cell culture conditions also included M-CSF, a critical growth and survival factor that induces a transcriptional program that promote macrophage differentiation, which is also essential for osteoclast maturation [65]. We compared mature and purified mononuclear and multinucleated macrophages, in order to minimize the combined effects of M-CSF and the lineage inducer (IFN-γ, IL-4, RANKL) and to study the pathways resulting from cell-cell fusion. We show that LGCs, FBGCs and osteoclasts are formed through cell-cell fusion, though we cannot exclude incomplete cytokinesis events [66]. The relative contribution of each event (incomplete cytokinesis vs. fusion) remains to be determined in human macrophages given that mouse MGCs form via modified cell division in granulomas [31].

We report that macrophage multinucleation causes a profound down-regulation of mononuclear phagocyte identity in LGCs, FBGCs and osteoclasts. This process consists of suppression of expression of major macrophage receptors (*CSF1R, MRC1, CD163, TLR2, CD74, IFNGR1*) and transcription factors (*MAFB, FOS, JUNB*). AP-1 transcription factors (*FOS, JUNB*) are involved in monocyte lineage specification [67, 68] and FOS is a well-described osteoclast lineage determinant transcription factor [28]. Given the role MAFB in tissue-resident macrophage lineage commitment [69], the results overall suggest a down-regulation of monocyte/macrophage identity caused by multinucleation. The intracellular signalling pathways leading to this ‘extinction’ phenotype remains to be determined. Since multinucleation of macrophages is associated with de-phosphorylation of kinases belonging to cJun NH2-terminal kinase (JNK) pathway such as Map3k1 [70], one can hypothesise the involvement of JNK and downstream transcriptional activity of AP-1 as a common suppressed signalling pathway associated with multinucleation in LGCs, FBGCs and osteoclasts. Similarly, the SRC kinase family member HCK is down-regulated in all three cell-types and dephosphorylated during macrophage multinucleation [70]. This suggests that signalling pathways crucial for macrophage migration, adhesion and function (Src, JNK, others) collectively contribute to the suppression of myeloid identity triggered by multinucleation. Multinucleation-induced suppression of core macrophage identity corroborates with previous findings in mice. First, the suppression of macrophage-specific transcripts such as Mafb and Csf1r have been shown in multinucleated mouse macrophages when compared to mononuclear cells cultured with M-CSF and a synthetic ligand for TLR2/6 [31]. Second, the downregulation of AP-1 was shown in BMDMs infected with Mtb [71]. Finally, work performed more than 50 years ago by Siamon Gordon and Zanvil A. Cohn on mouse macrophage-melanocyte heterokaryons and macrophage-macrophage homokaryons showed alteration of macrophage phagocytosis in fused cells, though the changes were more drastic in heterokaryons [72, 73]. In that sense, one limitation of our study is the lack of comparison between homologous and heterologous macrophage fusion since the latter can be relevant *in vivo* with the finding showing fusion of circulating blood monocytic cells with long-lived osteoclast syncytia in mice [15]. In cancer, the relevance of fusion between neoplastic cells and macrophages has been linked to tumour heterogeneity with acquired phenotypes in fused cells [74].

Macrophage multinucleation activates the lysosome-controlled cellular iron homeostasis in LGCs, FBGCs and osteoclasts. Our results suggest that multinucleation induces lysosomal biogenesis and capitalize on cytoplasmic and mitochondrial iron pools to sustain mitochondrial activity for increased energy demands. MGCs have been described having increased numbers of lysosomes decades ago [75] and several lines of evidence link v-ATPase-dependent lysosomal function to osteoclast multinucleation and function [76, 77]. Similarly, TFRC-mediated iron uptake induces mitochondrial respiration regulating osteoclast differentiation, mature osteoclast function and bone mass [78, 79]. Hence lysosomal biogenesis coupled with iron homeostasis is likely to be a ‘metabolic booster’ for specialized functions in multinucleated macrophages. Besides their degradative function critical in bone-resorbing osteoclasts, lysosomes can be seen as master regulatory organelles of energy homeostasis through iron dynamics and mitochondrial function [39-41]. Iron is essential for mitochondrial OXPHOs and TCA cycle activity in macrophages [80] and further studies on the lysosome-iron axis and its regulatory pathways will clarify the shared mechanistic insights of post-fusion metabolic reprogramming in MGCs.

Among MGC-specific transcriptomic signatures, the relatively enhanced phagocytic activity of multinucleated FBGCs and the increased spare respiratory capacity of multinucleated osteoclasts confirm the specialized functions of these MGCs. Mitochondrial function, and more specifically electron transport chain activity regulates osteoclast function [81, 82]. Our results show that cell-cell fusion and multinucleation enhance the mitochondrial activity required for resorptive activity. As previously reported, we found that, in addition to osteoclasts, MGCs are also TRAP+ [83] but among the three types of MGCs, only osteoclasts can have resorptive activity toward hydroxyapatite. FBGCs have been previously shown to have minimal resorptive activity but they cannot resorb bone [84]. Regarding the enhanced phagocytic activity of FBGCs when compared to osteoclasts and LGCs, the exact surface receptors used by FBGCs in phagocytosis of *S. Aureus* bioparticles remain to be identified. Given that all three types of MGCs show a downregulation of transcripts that belong to Fc receptors and complement C1Q family, the mechanisms of phagocytosis in these cells require further work. The specific upregulation of surface B7-H3 by LGCs suggest that this transmembrane protein is also involved in LGC-T cell crosstalk in granulomatous disease. B7-H3 mRNA is expressed by human mature osteoclasts [85] but our results showed that multinucleation of LGCs but not osteoclasts showed an up-regulation of membrane levels of B7-H3. Hence one explanation for the functional relevance of multinucleation in LGCs could be the spatial positioning of these MGCs expressing B7 family members such that they regulate T cells within granulomas. Altogether these results show drastic transcriptional and functional reprogramming caused by macrophage multinucleation. Whether these changes arise from all the nuclei or whether distinct functions are assigned to different nuclei of the MGC remain to be tested. Recent elegant work performed in multinucleated skeletal myofibers suggest transcriptional heterogeneity among the different nuclei of the polykaryon [86].

In summary, we showed that cell-cell fusion and multinucleation of LGCs, FBGCs and osteoclasts reset differentiated macrophages by at least two means: loss of monocyte/macrophage signature and triggering of lysosome-regulated intracellular iron metabolism. These common pathways confer cell type-specific gain of function that characterizes each polykaryon. We thus propose that cell fusion is a major determinant in shaping the context-dependent activity of MGCs.

## Materials and methods

### MGC generation from human donors

Human monocyte-derived macrophages were separated from healthy donor buffy coats by centrifugation through Histopaque 1077 (Sigma-Aldrich, Cat # H8889) gradient. Following Histopaque separation, peripheral blood mononuclear cells (PBMCs) were re-suspended in RPMI for osteoclasts (Life Technologies) and MEM-REGA 3 (1X) + 10% FBS (Life Technologies) for LGCs and FBGCs. For LGCs and FBGCs, monocytes were purified using a CD14 positive selection kit (StemCell Technologies, Cat # 17858). Monocytes (3.6 × 10^6^ cells) were seeded in 1 well Labtek chamber slides (ThermoFisher Scientific, Cat # 177372) and stimulated with 20 ng/ml M-CSF (PeproTech) for 3 days. At day 3, cells were differentiated with M-CSF (20 ng/ml, PeproTech) and IFN-γ (25 ng/ml, PeproTech, LGC differentiation) or IL-4 (10 ng/ml, PeproTech, FBGC differentiation) for further 4 days with the cell culture media changed every 2 days. For osteoclast generation, monocytes were purified by adherence for 1h at 37 °C, 5% CO_2_. The monolayer was washed 3 times with HBSS to remove non-adherent cells and monocytes were differentiated into macrophages for 3 days in RPMI media + 10% FBS containing 30 ng/ml M-CSF (PeproTech) and for further 4 days in media containing 30 ng/ml RANKL (PeproTech) and M-CSF. Studies on human donors were approved by Imperial College London and KU Leuven and have been performed in accordance with the ethical standards.

### Mononuclear and multinucleated macrophage sorting and purity

IFN-γ, IL-4 and RANKL-generated respective LGCs, FBGCs and osteoclasts were detached using Accutase (StemCell Technologies, Cat # 07920). Cells were labelled with 10 μg/ml Hoechst 33342 (Sigma-Aldrich, Cat # B2261) in cell sorting buffer containing PBS, 1% FBS (Sigma-Aldrich, Cat # F7524) and 2 mM EDTA for 40 min at 37°C to stain the DNA. Cells were then briefly centrifuged and re-suspended in ice-cold sorting buffer and kept on ice. Before FACS sorting, cells were filtered through a 70 μm nylon mesh (VWR, Cat # 732-2758). In order to sort mononuclear and multinucleated LGCs and FBGCs, FSC-A (logarithmic) versus Hoechst was plotted (**Supplementary Figure 1C**) and gating purity was further assessed by ImageStream (**Figure 1B, lower panel**). After sequential purity checks in at least 10 donors, a gating for mononuclear and multinucleated LGCs and FBGCs was selected (**See Supplementary Figure 1C**) based on 85-95% purity achieved by ImageStream in mononucleated and multinucleated cells. Cells were sorted on a BD FACSAria Fusion (BD Bioscience) using a 100 μm nozzle at a flow rate of 2000 events. Sorted cells were collected in FBS. For osteoclasts, after doublet exclusion, singlets with 1 nucleus (mononuclear cells) and more than 2 nuclei (multinucleated cells) were selected by FSC-A and SSC-A and the number of nuclei based on the Hoechst staining (**Supplementary Figure 1B**). Sorted mononuclear and multinucleated cells from each donor were run through the ImageStream that allowed direct visualisation of the number of nuclei; each cell population showed >95% purity.

### RNA extraction, library preparation and data analysis

Total RNA was extracted from sorted mononuclear and multinucleated LGCs, FBGCs and osteoclasts using Trizol (Invitrogen) and RNeasy mini kit (Qiagen) according to manufacturer’s instructions. Total RNA quality and concentration was analyzed using a NanoDrop 1000 spectrophotometer (ThermoFisher Scientific) and verified using Qubit meter (Invitrogen). Total RNA was analysed by Agilent 2100 Bioanalyzer (Agilent Tech Inc.) and RNA integrity number (RIN) values were ≥9.0 for all samples. Sequencing libraries for osteoclasts were prepared using NEBNext Ultra II Directional RNA Library Prep kit (Illumina). Briefly, RNA was purified and fragmented using poly-T oligo-attached magnetic beads using two rounds of purification followed by the first and second cDNA strand synthesis. Next, cDNA 3’ ends were adenylated and adapters ligated followed by 11 cycles of library amplification. The libraries were size selected using AMPure XP Beads (Beckman Coulter), purified and library quality was checked using Agilent 2100 Bioanalyzer. Samples were randomised to avoid batch effects and multiplexed libraries were run on a single lane (8 samples/lane) of the HiSeq 2500 platform (Illumina) to generate 100bp paired-end reads. Sequencing libraries for LGCs and FBGCs were prepared with the Lexogen QuantSeq 3’ mRNA-Seq Library prep kit according to the manufacturer’s instructions. Samples were indexed to allow for multiplexing. Library quality and size range was assessed using Agilent 2100 Bioanalyzer. Samples were randomised to avoid batch effects and multiplexed libraries were run on a single lane (8 samples/lane) of the HiSeq 4000 platform (Illumina) to generate 50bp single-end reads. Sequencing adapters were removed using Trimmomatic (v.0.36) and the reads quality was checked using FastQC (v.0.11.2) before and after trimming. Reads were aligned to the human genome (GRCh38.primary_assembly.genome.fa; annotation: gencode.v26.annotation.gtf) using HISAT2 package (v.2.1.0). Reads mapping to multiple loci in the reference genome were discarded. Mapping quality, read distribution, gene body coverage, GC content and rRNA contamination, were checked using picard (v.2.6.0) software. Gene level read counts were computed using HT-Seq-count (v2.7.14, annotation: gencode.v26.annotation.gtf). Genes with less than 10 aligned reads across all samples were filtered out as lowly expressed genes. Differential gene expression analysis between groups was performed using DESeq2 (v.1.14.1) and significantly differentially expressed genes were reported using fold-change at 1.5 times and below 5 % Benjamini-Hochberg (BH) adjusted p-value. In order to visualise the similarities between samples, unsupervised hierarchical clustering and principal component analysis (PCA) were performed using pcaExplorer (v.2.6.0, https://github.com/federicomarini/pcaExplorer) and pheatmap (v 1.0.10, https://CRAN.R-project.org/package=pheatmap) packages respectively. Volcano plots of differentially expressed genes were generated using enhanced volcano plot (v. 1.14.0, https://github.com/kevinblighe/EnhancedVolcano) package. All raw RNA-seq data processing steps were performed in Cx1 high-performance cluster computing environment, Imperial College London. Further analyses were conducted in R/Bioconductor environment v.3.4.4 (http://www.R-project.org/). Transcripts that are commonly down- or up-regulated were used for pathway analysis (EnrichR https://maayanlab.cloud/Enrichr/) with corresponding p-values (Fisher exact test) and adjusted p-values (Benjamini-Hochberg method).

### IncuCyte live cell imaging

At day 0, monocytes were resuspended in PBS and labelled with Incucyte Cytolight rapid green or red reagent for live cell cytoplasmic labelling (0.750 µM, Sartorius, Cat # 4705 and Cat # 4706) for 20 min at 37°C, 5% CO_2_ and washed with cold medium (MEM-REGA 3 (1X), Life Technologies). Labelled monocytes (0.5 × 10^6^ cells) were cultured at a ratio of red:green (1:1) in 8-well chamber slides (ThermoFischer Scientific, Cat # 154534) and stimulated with M-CSF (20 ng/ml, PeproTech, UK) for 3 days. Next, cells were stimulated with M-CSF (20 ng/ml, PeproTech, UK) in combination with IFN-γ (25 ng/ml, PeproTech, LGCs) or IL4 (10 ng/ml, PeproTech, FBGCs) or RANKL (30 ng/ml, PeproTech, osteoclasts). Imaging was performed with Incucyte live cell imaging system (Essen Biosciences) every 4 hours for the first 3 days and every 3 min after stimulation with either IFN-γ or IL-4 or RANKL with a magnification of 20X (37°C, 5% CO_2_). Phase images were acquired for every experiment and for fluorescent imaging, the acquisition times were 300 ms and 400 ms for green and red channels, respectively. The data are representative of 4 biological replicates.

### ImageStream analysis

At day 7 of MGC formation, differentiated LGCs, FBGCs and osteoclasts, were detached using Accutase (StemCell Technologies, Cat # 07920). Cells were stained with 10 μg/ml Hoechst 33342 (Sigma-Aldrich, Cat # B2261) in PBS supplemented with 1% FBS and 2 mM EDTA) for 40 min at 37°C to stain for DNA. Cells were then briefly centrifuged, re-suspended in ice-cold sorting buffer, incubated with FcR block (Miltenyi Biotec, Cat # 130059901) and stained with human anti-MRC1 or anti-B7-H3 (APC, Biolegend, MRC1, Cat # 321110 and B7-H3, Cat # 351006). Stained cells were filtered through a 70 μm nylon filter (VWR, Cat # 732-2758) and analysed with ImageStream (Amnis Corporation) with fluorescence measurements at 375 nm and 642 nm for Hoechst 33342 and MRC1/B7-H3, respectively. For each blood donor, a number of 50,000-100,000 cells were acquired. Based on their size (Area and Aspect Ratio) and number of nuclei, cells were divided into mononuclear and multinucleated cells (2+ nuclei) (Supplementary figure 2B). The mean fluorescence intensity of the specific markers was calculated and normalised by the mean area of the gated populations. Results were analysed with Ideas v5 Software (Amnis Corporation).

### Immunofluorescence

MGCs were culture in 8 well-plates (ThermoFisher Scientific, Cat # 177402) for 7 days. Cells were fixed in ice-cold methanol at -20 for 15 min, washed three times with PBS and blocked for 1h in PBS containing 5% normal goat serum (Cell signaling, Cat # #5425) and 0.3% Triton X-100. The cells were then incubated overnight at 4 °C with the primary antibody in blocking buffer. The antibodies used are as follows: rabbit monoclonal anti-CSF1R antibody (Cell Signalling Technologies, Cat # 67455S; 1:100), rabbit polyclonal anti-MRC1 antibody (Abcam, Cat # ab64693; 1:500). For visualisation, the secondary Alexa 455-conjugated anti-rabbit IgG (Cell signalling Technologies, Cat # 4413S; 1:500) was added to the blocking buffer for 1h at RT. Nuclei were stained in the Prolong Gold AntiFade Reagent with DAPI mounting medium (Cell Signalling Technologies, Cat # 8961S) and mounted under glass. Images were taken using epi-fluorescent Leica DM4B microscope and the raw fluorescence intensity was acquired using the ImageJ software 1.53. Multinucleated and mononuclear cells were visually identified based on their DAPI staining and multinucleated cells were delineated manually and their relative intensity of fluorescence (CSF1R and MRC1) was measured.

For CD3 immunofluorescence, prior to fixation, cells were washed two times with PBS to remove non-sticky cells. Next,cells were fixed in PBS, Hank’s balanced salt solution (HBSS Ca2+/ Mg 10X), BSA, sodium bicarbonate (NaHCO_3_) supplemented with 4% Paraformaldehyde (PFA, 40%, carl roth®) in 8-well plates (ThermoFisher Scientific, Cat # 177402). Cells were then washed twice and incubated 1h at RT in PFA (40%, carl roth®). Next, cells were rinsed thoroughly in HBSS buffer (PBS, HBSS Ca^2+^/ Mg 10X, BSA, NaHCO_3_) for 30 seconds and left to dry overnight at RT. Cells were then incubated 1h at RT in permeabilization buffer containing HBSS and 0.1% Triton-X. After washing, the fixed cells were incubated in blocking buffer (HBSS, FCS (10%), fragment crystallizable receptor (FcR) block (5%)) at RT for 2h. Following the blocking step, CD3-PE (Biolegend, Cat # 300456) was added and the cells were left for 2.5 hours in the dark. Hoechst 33342 (Sigma Aldrich) and Phalloidin (Invitrogen) were used to stain the nuclei and cytoskeleton, respectively. The plates were incubated for 1h at RT and covered from light. Thereafter, the slides were rinsed in HBSS buffer, dried and covered with few drops of diamond mounting medium (Invitrogen, Cat # P36961) were added before microscopy analysis (20X magnification, Zeiss Axiovert 200M, Carl Zeiss).

### DC-STAMP siRNA transfection

siRNA transfection of primary human macrophages was performed based on the previously published protocols [80, 87]. Monocytes (3.6 ×10^6^ cells) were first seeded and stimulated with M-CSF for 3 days (20 ng/ml, PeproTech) in 1-well chamber slides (ThermoFisher Scientific, Cat # 177372) (72 h, 37°C, 5% CO_2_). Macrophages were then transfected using Dharmafect 1 (Dharmacon, Cat # T-2001-03) diluted in OPTIMEM medium (1:50, Invitrogen) with human *DC-STAMP* siRNA (100 nM, Dharmacon) and non-targeting scrambled siRNA used as a control. After 8 hours, cells were washed and differentiated into MGCs with stimulatuion with either IFN-γ (25 ng/ml, PeproTech) or IL-4 (10 ng/ml, PeproTech) or RANKL (30 ng/ml, PeproTech) in presence of M-CSF (20 ng/ml, PeproTech) containing media for 2 days. The effect of *DC-STAMP* knockdown was subsequently measured by quantitative RT-PCR (qRT-PCR) in LGCs and FBGCs (Supplementary Figure 2E). For osteoclasts, DC-STAMP si-RNA transfection was performed in Pereira et al. and the cDNA was used for the described qPCR measurements [21].

### Quantitative RT-PCR

Total RNA was extracted using the Trizol reagent (Invitrogen) according to the manufacturer’s instructions. For osteoclasts, complementary DNA (cDNA) was synthesised using iScript cDNA Synthesis Kit (Bio-Rad). A total of 10 ng cDNA for each sample was used and all qRT-PCR reactions were performed on a ViaA 7 Real-Time PCR System (Life Technologies) using Brilliant II SYBR Green QPCR Master Mix (Agilent). Results were analysed by the comparative Ct method using ViiA 7 RUO software, and each sample was normalised relative to *HPRT* expression. For LGCs and FBGCs, complementary DNA (cDNA) was prepared using Superscript II reverse transcriptase (Invitrogen, Cat # 4311235) and random primers (Invitrogen, Cat # 4319979). qRT-PCR was performed using a TaqMan gene expression assay (Applied Biosystems) on a 7500 Real-Time PCR System Apparatus. Results were analysed by the comparative Ct method and each sample was normalised to *HPRT* expression levels. Primers used are listed in **Supplementary Table 7**.

### Lysosomal function, iron supplementation and MGC fusion

Monocytes were seeded and stimulated with M-CSF (20 ng/ml, PeproTech) in 8-well chamber slides (ThermoFisher Scientific, Cat # 177402) for 3 days (37°C, 5% CO_2_). Cells were the re-incubated with M-CSF (20 ng/ml, PeproTech) in combination with either IFN-γ (25 ng/ml, PeproTech) or IL-4 (10 ng/ml, PeproTech) or RANKL (30 ng/ml, PeproTech). Simultaneously, cells were incubated with either Bafilomycin A1 (Baf A1, Sanbio B.V., Cat # 11038-500), Concanamycin A (Con A, Sigma-Aldrich, Cat # 27689), Hydroxychloroquine sulfate (Hydroxy Sulfate, Sigma-Aldrich, Cat # 379409) and Ammonium chloride (NH4Cl, Sigma-Aldrich, Cat # A9434). As part of the rescue experiment, iron chloride (FeCl_3,_ Sigma-Aldrich, Cat # 451649) was added to some cells at day 4 and cells were incubated for further 2 days at 37°C, 5% O_2_, after which Giemsa staining was performed to measure fusion index using ImageJ. Fusion index refers to a specific area and equals to the percentage of [number of nuclei (>3)] / [total number of nuclei].

### Phagocytosis by ImageStream

Differentiated LGCs, FBGCs and osteoclasts were detached using Accutase (StemCell Technologies, Cat # 07920) and were filtered through a 70 µm nylon mesh (VWR, Cat # 732-2758). For DNA staining, cells were labelled with 10 µg/ml Hoechst 33342 (Sigma-Aldrich, Cat # B2261) in PBS supplemented with 1% FBS and 2 mM EDTA. Cells were then centrifuged and re-suspended in cold PBS. Phrodo red *S. Aureus* bioparticles (Invitrogen, Cat # A10010) were added according to the manufacturer’s instructions and incubated for 30 min at 37 °C. Cells were re-centrifuged, fixed in 0.4% formaldehyde and analysed with ImageStream (Amnis Corporation). Fluorescence was measured by ImageStream at 375 nm (Hoechst) and 581 nm (Phrodo red *S. Aureus* bioparticles). Based on their size and Hoechst staining, cells were categorised as mononuclear and multinucleated cells. The mean fluorescence of Phrodo red *S. Aureus* bioparticles was measured as a readout of phagocytic activity of the cells. Results were analysed with Ideas v5 Software (Amnis Corporation).

### Granuloma-like cluster formation

To generate granuloma-like cell clusters, cell differentiation was achieved by using PBMCs instead of isolated monocytes from blood donors. PBMCs (0.7 × 10^6^) were incubated with conditioned media (MEM-REGA 3 (1X) + 10% FBS (Life Technologies)) containing M-CSF (20 ng/ml, PeproTech) and 10% FBS (Sigma-Aldrich, Cat # F7524) for 3 days. At day 3, cells were supplemented with either IFN-γ (25 ng/ml, PeproTech) or IL-4 (10 ng/ml, PeproTech) or RANKL (30 ng/ml, PeproTech). At day 7 of differentiation, cell culture media was removed and cells were washed twice in PBS prior to Giemsa staining and immunofluorescence analysis.

### Extracellular flux analysis

Real-time measurements of oxygen consumption rate (OCR) and extracellular acidification rate (ECAR) were performed using a Seahorse XF96 Extracellular Flux Analyzer (Agilent Technologies). Donor-derived human osteoclasts were cultured for 7 days and sorted as mononuclear and multinuclear cells (see description above). Cells were then washed and 5 × 10^5^ osteoclasts were seeded onto a XF96 plate containing Seahorse XF RPMI medium (Agilent Technologies). The cells were left for 1 h at 37 °C after which the different metabolic drugs were injected (oligomycin 1μM, FCCP 2μM, rotenone/antimycin 1μM, 2-DG 50mM) during real-time measurements of OCR and ECAR, using the Seahorse XF Cell Mito Stress Test Kit (Agilent Technologies). Basal respiration was calculated as [the last measurement before addition of oligomycin – non-mitochondrial respiration (minimum rate measurement after Rot/AntA)]. Maximal respiration is shown as [the maximum rate measurement after addition of FCCP – non-mitochondrial respiration]. Estimated ATP production designates [the last measurement before addition of oligomycin – minimum rate after oligomycin]. Glycolysis refers to ECAR values before the addition of oligomycin.

### Mitochondrial copy number

Total DNA was extracted from cell samples using the DNeasy Blood and Tissue kit (Qiagen, Valencia, CA, USA) with proteinase K and RNase treatment, according to the manufacturer’s instructions. To quantify mtDNA copy number, real-time qPCR was performed using the ViiA 7 (Life Technologies) to measure gene expression against external standards for mitochondrial (tRNA Leu) and nuclear DNA (β2-microglobulin), as previously described [88]. Mitochondrial copy number was calculated using the formula (mt copy number = 2*2^(nuclear copy number – mitochondrial copy number)) for mononuclear and multinuclear macrophage population for each donor. The quantification of mitochondrial transcripts in Figure 6F was then normalised by the mitochondria copy number (Figure 6E) to account for mitochondria numbers.

### Bone resorption assay

The resorption activity of human osteoclasts was measured *in vitro* using Osteo Assay Surface 96-well plates (Corning) which are made of hydroxyapatite surfaces. MGCs were cultured until day 4 and incubated with cell dissociation buffer (Sigma; Catalogue # C5914). A total of 10^5^ cells/well were seeded onto Osteo Assay Surface Plates. After 2 days of culture with the MGC-specific cytokines, the wells were rinsed twice in PBS and incubated with 10% bleach solution for 30 min at RT. The wells were then washed twice in PBS and allowed to dry at room temperature. Individual resorption pits were imaged by light microscopy. Images were inverted and processed using Photoshop to yield high-contrast images and show the resorbed areas in black against a white background. Binary images of each individual well were then subjected to automated analysis (ImageJ), using constant ‘threshold’ and ‘minimum particle’ levels, to determine the number and surface area of resorbed pits.

### Giemsa and TRAP staining

Cells were fixed with methanol for 5 seconds and stained with Giemsa (1:10 dilution with MiliQ, VWR) for 17 min. After staining, cells were washed two times with water. For TRAP staining, cells were stained for tartrate-resistant acid phosphatase (TRAP), as previously described [89]. TRAP+ multinucleated cells (more than three nuclei) were defined as MGCs.

### Statistical analysis

Data were analysed using GraphPad Prism software, version v.9.0.1 (GraphPad Software). Unless indicated, graphs show mean +/- SD for each group and the statistical test is indicated in the corresponding legend. Fusion index data (Figure 3E) were log transformed and significance was tested by one-way ANOVA followed by Sídàk’s multiple comparison test. Significance is based upon a P-value less than 0.05.

## Acknowledgements

We would like to thank Carla Rios Luci from the NanoHealth and Optical Imaging Group for her support with the ImageStream experiments. Illustrative cartoons were designed with BioRender.com. This research paper was funded by the Medical Research Council (UK) and Fonds Wetenschappelijk Onderzoek Vlaanderen (Research Foundation Flanders, FWO-Vlaanderen).

## FIGURE LEGENDS

**Supplementary Figure 1.**
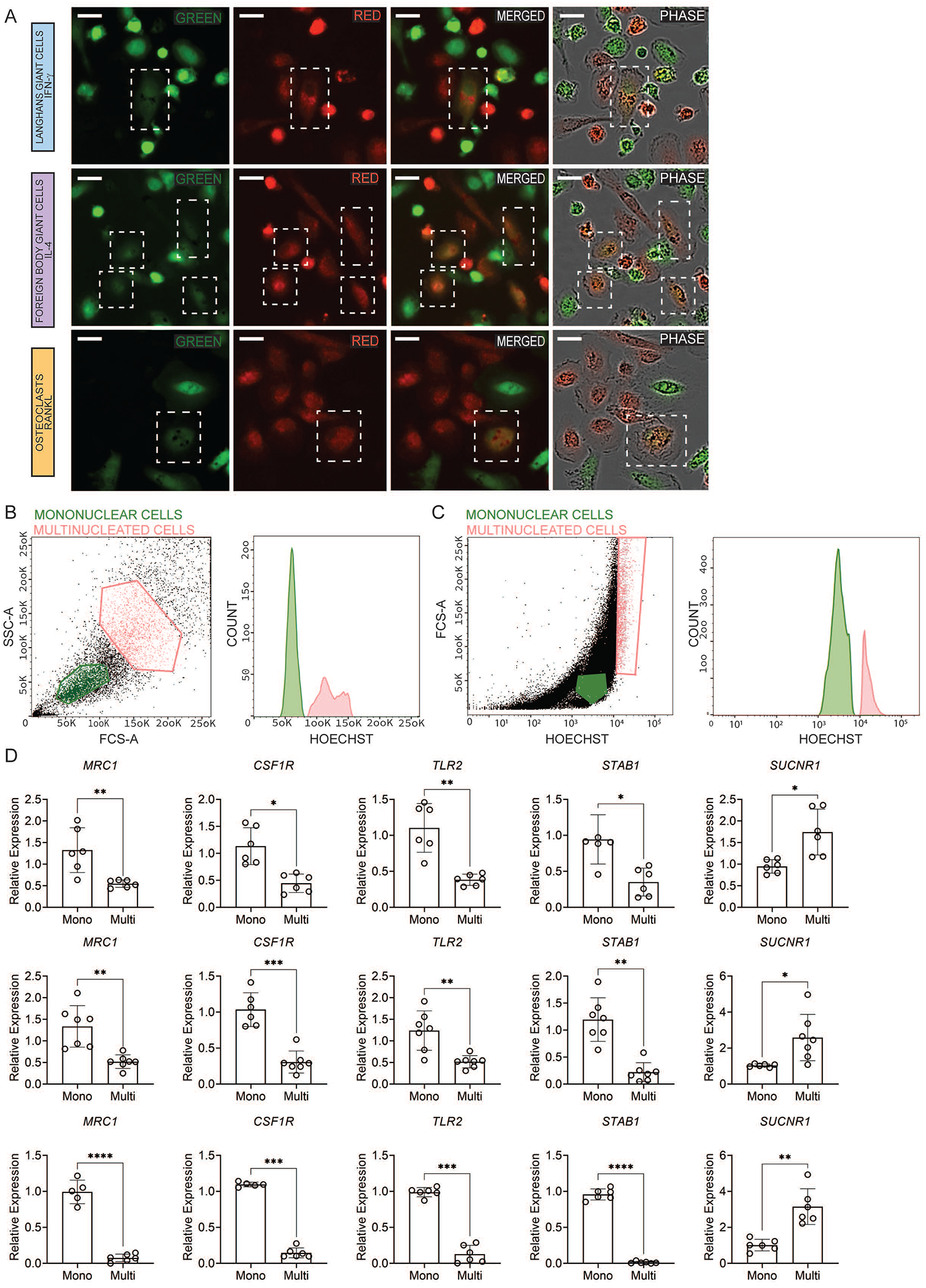
Multinucleated giant cells are generated through cell-cell fusion. **(A)** Isolated monocytes were labelled with either green or red dyes and stimulated with IFN-γ (LGCs), IL-4 (FBGC), or RANKL (osteoclasts) for giant cell formation. Cell-cell fusion was monitored with live cell imaging system (Incucyte). Orange dye-labelled giant cells (fusion between red and green) are shown within dotted white boxes. **(B)** FACS sorting strategy for mononuclear and multinucleated (> 2 nuclei) osteoclasts based on size and Hoechst (DNA content as a readout of multinucleation). **(C)** FACS sorting strategy for mononuclear and multinucleated (> 2 nuclei) LGCs and FBGCs based on size and Hoechst (DNA content as a readout of multinucleation). **(D)** *MRC1, CSF1R, TLR2, STAB1 and SUCNR1* relative expression measured by qRT-PCR in LGCs (upper), FBGCs (middle) and osteoclasts (lower), in sorted mononuclear and multinucleated cells; at least n=6 donors. Error bars are mean ± SD; significance tested by paired t-test; *, P < 0.05; **, P < 0.01; ***, P<0.001; ****, P < 0.0001. Scale bar, 200μm **(A)**.

**Supplementary Figure 2.**
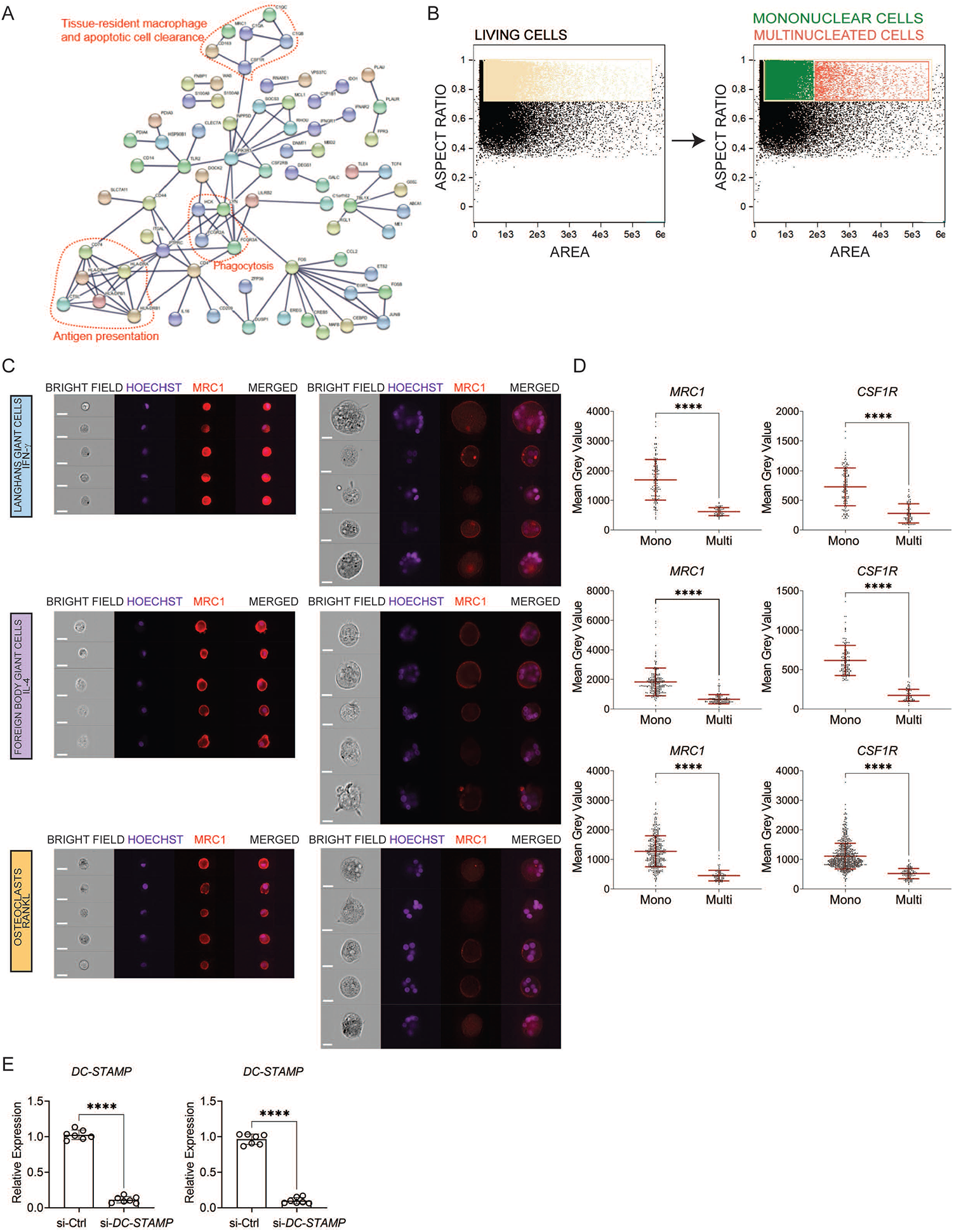
Fusion and multinucleation causes the down-regulation of a shared mononuclear phagocyte gene signature between LGCs, FBGCs and osteoclasts. **(A)** Protein– protein interaction (PPI) network in 191 commonly down-regulated genes in LGCs, FBGCs and osteoclasts illustrated by STRING (high confidence score=0.9, only connected nodes are shown). Dashed lines denote the genes belonging to antigen presentation, phagocytosis and tissue resident macrophage and apoptotic cell clearance pathways. **(B)** ImageStream gating for mononuclear and multinucleated cells shown in LGCs. The gating strategy for FBGCs and osteoclasts is similar. **(C)** ImageStream showing bright field, nuclei staining (Hoechst), and MRC1 (red) staining in mononuclear and multinucleated LGCs, FBGCs and osteoclasts. **(D)** MRC1 and CSF1R immunofluorescence quantification in mononuclear and multinucleated LGCs, FBGCs and osteoclasts; n=2 donors. **(E)** DC-STAMP expression following its knockdown in LGCs (left) and FBGCs (right). si-Ctrl, scrambled siRNA; si-DC-STAMP, DC-STAMP siRNA; n=7 donors. Error bars are mean ± SD; significance tested by unpaired (**D**) and paired (**E**) t-test; ****, P < 0.0001; scale bar, 20µm **(C)**.

**Supplementary Figure 3.**
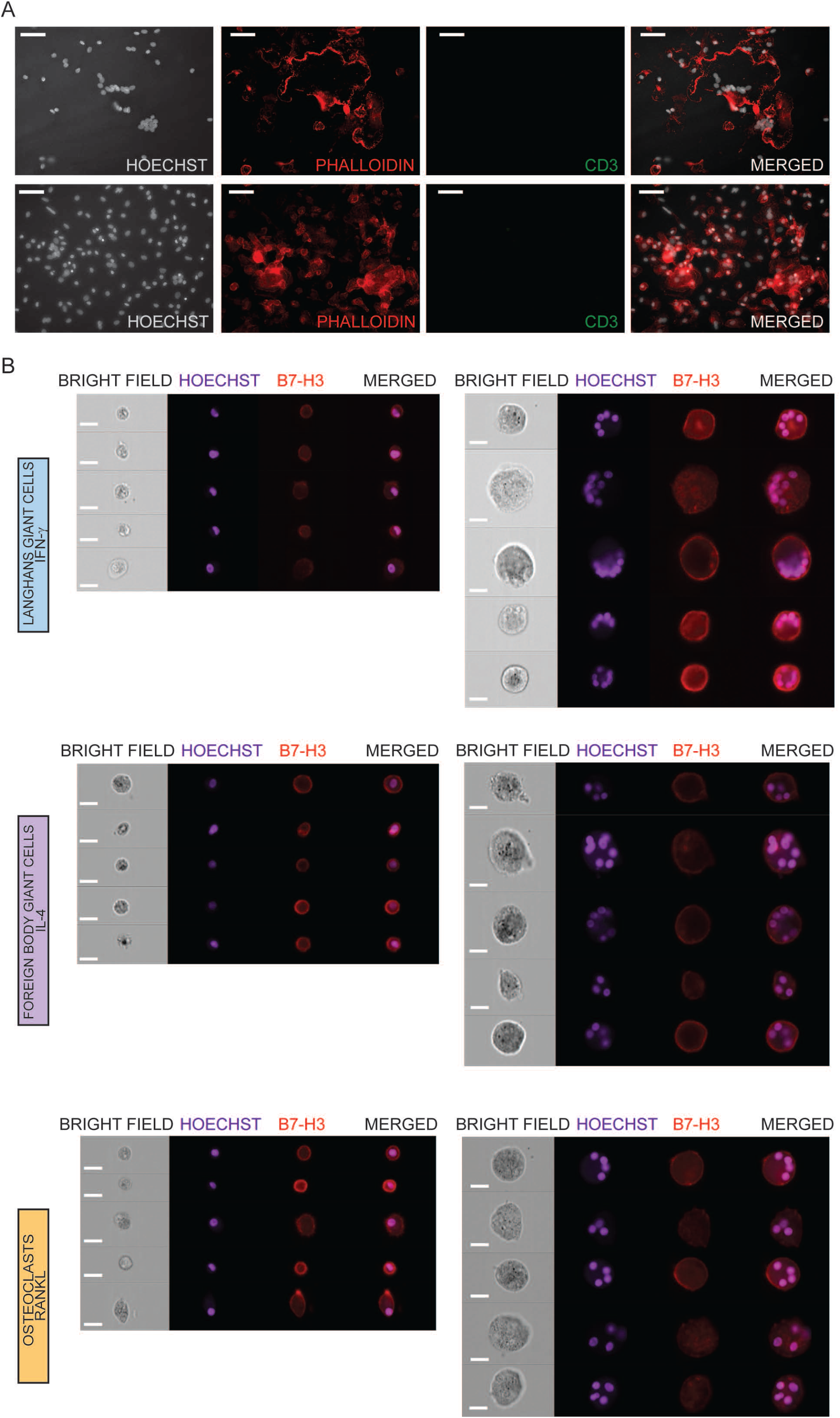
**(A)** CD3 immunofluorescence (green) in PBMC-derived FBGCs (upper panel) and osteoclasts (lower panel). Hoechst (grey) and phalloidin (red) staining show the nuclei and cytoskeleton, respectively. **(B)** LGCs show increased membrane expression of B7-H3. ImageStream showing bright field, nuclei staining (Hoechst), and B7-H3 (red) staining in mononuclear and multinucleated LGCs, FBGCs and osteoclasts. Scale bar, 20µm.

**Supplementary Figure 4.**
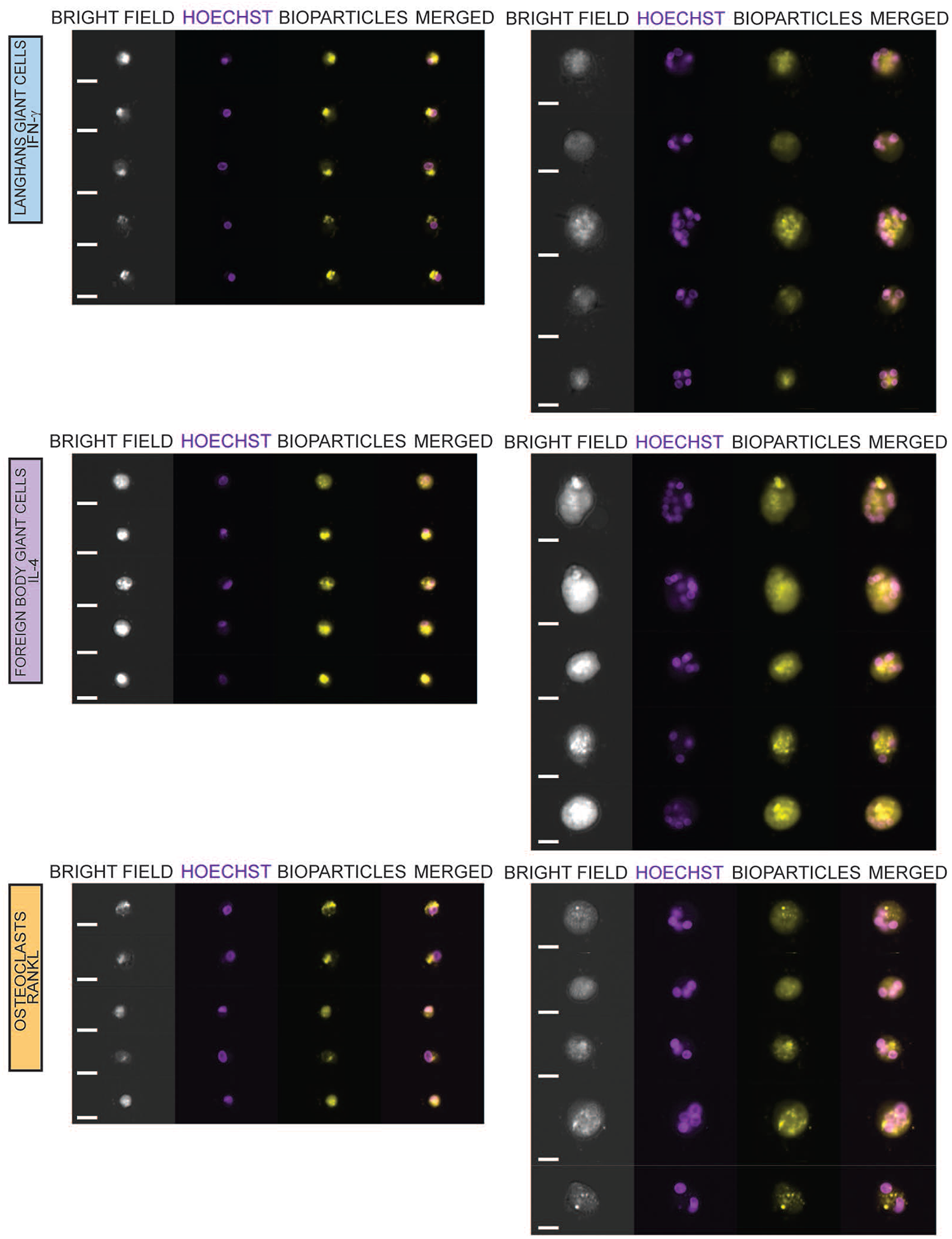
FBGCs show a distinctively enhanced phagocytic capacity. ImageStream showing bright field, nuclei staining (Hoechst), and *S. Aureus*-coated bioparticles (yellow) staining in mononuclear and multinucleated LGCs, FBGCs and osteoclasts. Scale bar, 20µm.

